# 3D Reconstruction of Nanoparticle Distribution in Tumor Spheroids with Volume Electron Microscopy

**DOI:** 10.64898/2026.04.17.719153

**Authors:** Davide Bottone, Lukas R. H. Gerken, Sebastian Habermann, José María Mateos, Miriam S. Lucas, Johannes Riemann, Melanie Fachet, Ute Resch-Genger, Vera M. Kissling, Matthias Rösslein, Alexander Gogos, Inge K. Herrmann

## Abstract

Spatially resolved characterization of nanomaterial (NM) distribution within cellular ultrastructure is essential for understanding NM fate and activity in biological systems. Volume electron microscopy (vEM) is uniquely positioned to address this challenge, yet fully documented quantitative pipelines that simultaneously segment NMs and cellular structures remain scarce. Here, an end-to-end analytical pipeline is presented based on the example of serial block-face scanning electron microscopy (SBF-SEM) data of tumor spheroids containing nanoparticles (NPs). A hybrid segmentation strategy is adopted: a fine-tuned Cellpose-SAM model for cells and nuclei, and an empirical Bayes approach for AuNPs. The fine-tuned model outperforms both the pre-trained baseline and benchmark experiments in Amira, and shows good generalization to 2D EM datasets of varying sample types, suggesting potential as a general-purpose segmentation model for electron microscopy. Full 3D reconstruction of NP distributions reveals preferential clustering in the perinuclear region, with a median nucleus-to-NP distance of 2.57 µm and NM uptake spanning several orders of magnitude across cells. Furthermore, morphological analysis of segmented cells and nuclei using 3D shape descriptors and local curvature metrics provides quantitative access to features inaccessible from single sections. Together, these results establish a reproducible, open framework for the joint quantitative analysis of NM distribution and cellular morphology in vEM data.

## 1 Introduction

Nanomaterials (NMs) have emerged as versatile platforms for biomedical applications, with tunable physicochemical properties enabling control over their interactions with biological systems at the molecular and cellular level [1–3]. However, successful clinical translation critically depends on a detailed understanding of cellular uptake, intracellular localization, and NM fate [4–6], requiring analytical approaches that resolve NM distributions within the full context of cellular ultrastructure.

Electron microscopy (EM) occupies a unique position in this regard due to its ability to visualize biological ultrastructure and electron-dense NMs such as gold nanoparticles (AuNPs) at nanometer-scale resolution. Recent advances in volumetric EM [7, 8], particularly serial block-face scanning electron microscopy (SBF-SEM) [9] and focused ion beam SEM (FIB-SEM) [10], now allow three-dimensional (3D) reconstruction of cellular architectures and NM distributions simultaneously. Yet, extracting quantitative information from these datasets remains a major computational challenge [11–13], as volumetric analysis relies on accurate segmentation of structures of interest within large, heterogeneous image stacks, a task that has historically been manual and difficult to scale.

In recent years, deep learning methods have begun to address this bottleneck. Generalist frameworks such as Cellpose [14] can delineate cells and nuclei across heterogeneous microscopy datasets without extensive retraining. Cellpose-SAM [15] extends this by leveraging the image encoder of the Segment Anything Model (SAM) [16] and predicting Cellpose vector flow fields directly from encoder outputs. However, both frameworks were developed and benchmarked on light microscopy data, making generalization to EM image characteristics a non-trivial challenge. Furthermore, despite the progress in deep learning-based image analysis, reproducibility remains a concern [17–19], and the field still lacks studies that provide simultaneously: fully documented volumetric datasets, transparent analysis pipelines, and systematic benchmarking across imaging conditions [20]. This is particularly true for vEM datasets, which are less abundant and typically harder to characterize than other volumetric microscopy techniques [8].

Crucially, only a small number of works have performed quantitative 3D analysis of vEM datasets that segment NMs together with their surrounding cellular ultrastructure [21, 22]. Most studies focus on either the NMs [23, 24] or the cellular compartments in isolation, or rely on correlative strategies that combine fluorescence labeling with vEM [25] or cryo-X-ray tomography [26] rather than direct segmentation within a single volumetric dataset. This limits the ability to quantify NM spatial distributions, clustering, and cell-to-cell heterogeneity in uptake, a well-documented but mechanistically complex phenomenon [27, 28], in relation to precise structural context. Beyond NM localization, 3D segmentation of cells and nuclei opens access to morphological information that is inaccessible from single sections, with 3D shape descriptors providing quantitative readouts of the cellular state [29, 30].

Here, we address these challenges using a tumor spheroid model [31], a 3D platform recapitulating key features of the tumor microenvironment, in which we simultaneously segment cells, nuclei, and AuNPs from vEM data to quantify NP uptake and intracellular distribution, as well as global and local shape descriptors for cells and nuclei. We present a quantitative analysis pipeline built on a refined Cellpose-SAM segmentation model, release the underlying annotated dataset as a public resource, and benchmark our approach against the AI-assisted segmentation tool in Amira 3D software suite, demonstrating comparable accuracy. We further show that the pipeline generalizes to datasets from different EM platforms and sample types. Simultaneous 3D segmentation of AuNPs and cellular compartments enables quantitative morphological characterization of cells and nuclei alongside spatially resolved NP uptake analysis, providing a scalable framework for structure-function studies in nanomedicine.

## 2 Materials and methods

### 2.1 Spheroid preparation

FaDu (HTB-43, ATCC) cell spheroids were prepared according to the protocol reported by Gerken et al. [31]. Briefly, 40 000 cells were seeded in 1.15 mL growth medium into 12-well plates and allowed to adhere for 24 h at 37 °C in a humidified incubator with 5 % CO_2_. Then, the medium was exchanged with fresh one with a 50 µg mL^−1^ Au NPs concentration, which was obtained by adding 57.5 µL of a 1 mg mL^−1^ Au NPs suspension (BioPure Gold Nanospheres, nanoComposix; citrate-capped; 50 nm ±4 nm diameter, used as received) to 1.092 mL of growth medium. After 24 h NP exposure, the cells were detached using trypsin, washed (i.e. the medium removed and replaced with NP-free medium) and transferred to ultra-low attachment plates for spheroid formation [31], with an initial cell seeding density of 500 cells per well. Additionally, the spheroid growth medium was spiked with Au NPs (50 µg mL^−1^) to enable continued exposure of the growing spheroid. After 24 h, the medium was exchanged with fresh medium without NPs. On day 2 after spheroid seeding, spheroids were harvested and fixed with 2.5% glutaraldehyde (EM Grade, Sigma-Aldrich) in 0.1 mol L^−1^ sodium cacodylate buffer (Electron Microscopy Sciences, pH 7.4), and stored in fixative at 4 °C for several days before additional processing.

The samples were then stained and embedded in resin according to a reduced osmium–thiocarbohydrazide–osmium protocol with additional uranyl acetate and lead aspartate contrasting, followed by dehydration and embedding in Epon–Araldite resin [32, 33]. The full details of the protocol can be found in the Supplementary Methods (Section S1.1).

Light microscopy images of the spheroids were recorded before harvesting for estimation of their size prior to resin embedding.

### 2.2 vEM data acquisition

The EM volume dataset was acquired with a serial block-face scanning electron microscope (SBF-SEM, Thermo Fisher Apreo Volumescope) with an *XY Z* voxel size of 10 nm ×10 nm ×50 nm, using a T1 segmented lower in-lens detector in A+B with accelerating voltage 1177.33 V, dwell time 0.5 µs, working distance 6.258 mm, and beam current 100 pA. Images were acquired with a 102.4 × µm 102.4 µm field of view at the center of a single spheroid, which was sectioned for 35 µm. Images containing acquisition or sectioning errors were manually excluded from the analysis.

### 2.3 Denoising

Images were denoised with Noise2Void [34, 35] in the CAREamics implementation (https://github.com/CAREamics). The model’s UNet architecture [36] was configured with 32 initial channels, depth 2, and 64 × 64 patch size. The model was trained for 100 epochs on the first 10 slices of the volume stack, with an 80/20 training/validation split, using the default CAREamics training options (Adam optimizer [37], 0.001 learning rate, learning rate reduction on plateau). Each *XY* slice of the volume stack was denoised separately, and intensities were rescaled per-slice in the 0-255 range.

### 2.4 Registration

The denoised images were registered using a custom pair-wise intensity-based approach implemented in Pytorch [38]. An affine transform with full 2D degrees of freedom was learned per image pair by minimizing the loss ℒ_pair_ = 1− ρ_NCC_, where ρ_NCC_ is the normalized cross correlation between the two images. Optimization was carried out sequentially on a 4-level multi-resolution pyramid, from coarse to fine, with downscaling factors (8, 4, 2, 1) and learning rates (1 ×10^−3^, 1 ×10^−3^, 5 ×10^−4^, 1 ×10^−4^)for an Adam optimizer [37]. Each level of the pyramid was optimized for 300 epochs, using an early stopping criterion with a tolerance of 1× 10^−5^ and patience 15. The registered volume stack was then automatically cropped to remove empty borders introduced by image warping.

#### 2.4.1 Segmentation of cellular structures

Ground truth segmentation masks for cellular structures (cells, nuclei) were manually recorded from the denoising images with the help of the microSAM plugin [16, 39] in napari [40]. A total of 415 cells and 212 nuclei were labeled across 10 randomly selected 2D *XY* images. Additionally, 10 vertical slices (5 *X Z*, 5 *Y Z*) were randomly selected from the registered volume stack after resampling to isotropic voxel spacing (50 nm ×50 nm ×50 nm) to provide additional labeled data covering all three spatial orientations; 215 cells and 92 nuclei were labeled across these slices. In total, the labeled dataset comprises 630 cells and 304 nuclei across 20 images; note that these counts are not of unique objects, as the same cell or nucleus may appear in multiple slices of the same volume.

Cellpose-SAM [14, 15] was fine-tuned using all labeled data (*XY, X Z*, and *Y Z* slices) via the provided API. The images were resampled to (100 nm ×100 nm ×100 nm) spacing prior to training, and split with an 80/20 ratio between training and validation. Two separate models were trained, targeting respectively cells and nuclei, with learning rate 5 ×10^−5^ and batch size 8. The cells model was trained for 1000 epochs, while the nuclei model was trained for 500 epochs.

Prediction on the full volume was then run at the resampled (100 nm ×100 nm ×100 nm) resolution. Before mask generation, a median filter of size 3 was applied to the predicted probability logits. For nuclei, the predicted flows showed several artifacts that can be represented as a non-zero curl component. Such artifacts arise from the more complex shapes of nuclei compared to cells, with pronounced concavities resulting in competing flow directions. Moreover, the ground truth flow field of Cellpose is by construction irrotational, being the gradient of a scalar potential [14]. The calculated flows **V** can be decomposed into irrotational (curl-free) and solenoidal (divergence-free) components using the Helmholtz-Hodge decomposition [41], as **V** = **V**_ir_ +**V**_**sol**_. The irrotational flow component was calculated in the Fourier domain as [42]:

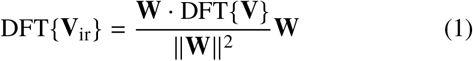

where DFT {·} denotes the discrete Fourier transform operator, and **W** are the finite-difference wave vectors at frequency ω. DFT was calculated using Fast Fourier Transform (FFT), and the irrotational flow component was calculated by applying the inverse transform to the result of Equation 1.

The processed probability logits and flows were then used for mask generation using the Cellpose API, with a probability threshold of −0.5 and a minimum diameter of 4.5 µm for cells and 3.7 µm for nuclei. Oversegmented nuclei masks were merged using a region adjacency graph (RAG) approach. Graph edge weights *w*were calculated as *w* = 1− *w*_prob_ +*w*_edge_, where *w*_prob_ is the probability corresponding to the average cell logit at label boundary, and *w*_edge_ is the average normalized image edges, calculated using a Gaussian edge map with σ = 2. Lower values of *w* are typically indicative of oversegmentation. Label contact surface was calculated with a voxel-shift approach, and the relative contact surface between two labels *A*_rel_ = *A*/*A*_tot_ was added as an additional graph edge weight. The RAG was merged with conditions *w* ≤ 0.15 and *A*_rel_ ≥ 0.15.

The final masks were upsampled at (50 nm ×50 nm ×50 nm) resolution via geodesic dilation. Seeds were obtained via nearest-neighbor interpolation of the low-resolution masks, followed by two consecutive erosion steps. The foreground constraint was defined by thresholding the linearly interpolated probability logits at 0.

Instance segmentation accuracy was evaluated by calculating average precision AP_*τ*_ at different values τ of the Jaccard index IoU. A predicted instance *P*_*i*_ and ground truth instance *G* _*j*_ were matched if:

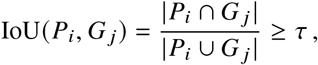

where *P*_*i*_ ∈ 𝒫 and *G* _*j*_ ∈ 𝒢 are individual instances from the predicted and ground truth sets, respectively, and | · | denotes pixel count. Using one-to-one matching, we calculated true positive TP_*τ*_, false positive FP_*τ*_, and false negative FN_*τ*_ counts. Average precision was then calculated as:

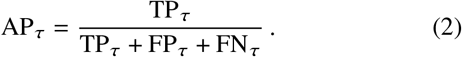

Additionally, voxel-level instance delineation accuracy was evaluated with aggregated Jaccard index (AJI) [43] in the Cellpose implementation. Segmentation accuracy was evaluated on the full set of labeled slices. It should be noted that the labeled 2D slices were also used for model fine-tuning; however, the evaluated output is that of the complete 3D pipeline, which runs independent 2D predictions across all three spatial orientations and combines them through flow integration, merging, and upsampling. The 3D predictions are therefore substantially different from a direct reproduction of the 2D training labels, and the reported metrics reflect the performance of the end-to-end pipeline rather than 2D model generalization alone.

The fine-tuned Cellpose-SAM models were additionally bench-marked against the AI-assisted segmentation tool in Amira (Thermo Fisher Scientific). For this task, a human-in-the-loop workflow based on a UNet [36] architecture with a VGG19 [44] backbone was employed, and separate models were trained for cells and nuclei. A detailed description of the training procedure can be found in Section S1.2.

### 2.5 Generalization to 2D SEM datasets

Various 2D SEM datasets of cell cultures spiked with NPs were used to assess qualitatively the generalization of the fine-tuned Cellpose-SAM models for cell segmentation, and are summarized in Table 1. The FaDu-Au [31] and MT-Ti-MIL [46] datasets were previously published by our group, while the HT1080-Hf dataset was acquired with a new imaging campaign conducted on previously published samples [45].

**Table 1:**
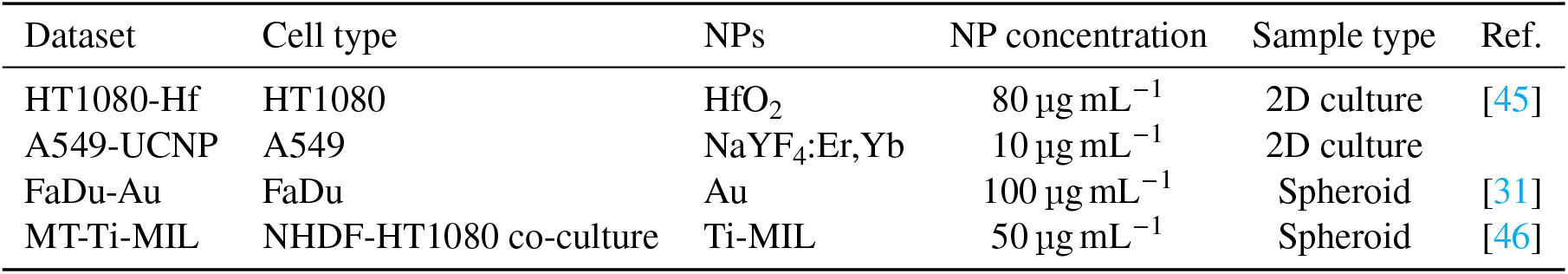
2D SEM datasets used for assessing the generalization of the fine-tuned Cellpose-SAM models for cell segmentation.

| Dataset | Cell type | NPs | NP concentration | Sample type | Ref. |
| --- | --- | --- | --- | --- | --- |
| HT1080-Hf | HT1080 | HfO<sub>2</sub> | 80 $\mu\text{g mL}^{-1}$ | 2D culture | [45] |
| A549-UCNP | A549 | NaYF<sub>4</sub>:Er,Yb | 10 $\mu\text{g mL}^{-1}$ | 2D culture | |
| FaDu-Au | FaDu | Au | 100 $\mu\text{g mL}^{-1}$ | Spheroid | [31] |
| MT-Ti-MIL | NHDF-HT1080 co-culture | Ti-MIL | 50 $\mu\text{g mL}^{-1}$ | Spheroid | [46] |

The A549-UCNP dataset, on the other hand, was prepared for this work. Briefly, human lung adenocarcinoma A549 cells were maintained in Dulbecco’s Modified Eagle Medium (DMEM) supplemented with 10 % fetal bovine serum at 37 °C in a humidified incubator with 5 % CO_2_. For NP exposure, A549 cells were seeded in T75 flasks and grown until a confluence of 70 % to 80 %. The cells were spiked for 24 h under standard culture conditions at a final concentration of 10 µg mL^−1^ with oleate-capped luminescent lanthanide-based NaYF_4_:Er,Yb upconversion nanoparticles (UCNPs) synthesized as described here [47]. After exposure, cells were washed three times with 0.1 mol L^−1^ HEPES buffer to remove unbound nanoparticles and fixed immediately using 4 % paraformaldehyde in 0.1 mol L^−1^ HEPES buffer for 1 h at room temperature. After overnight fixation, cells were centrifuged to form a pellet (200 g, 5 min) and were washed three times with 0.1 mol L^−1^ sodium cacodylate buffer for 3 min. To visualize cellular structures, our previously published protocols were followed [46, 48, 49]. In brief, the cell pellets were stained for 1 h at room temperature with 1 % OsO_4_ (Electron Microscopy Sciences) in 0.1 mol L^−1^ cacodylate buffer. Thereafter, three washing steps with Millipore water for 3 min each were performed. The cell pellets were then dehydrated through the following ethanol series: 30 % (5 min), 50 % (5 min), 70 % (5 min, 90 % (5 min), and 100 % (3 ×10 min). Overnight, the cell pellets were incubated with a 1:1 mixture of 100 % Epon (Epon 812 substitute resin, Epoxy embedding kit 45359, Sigma-Aldrich) and 100 % ethanol. The cell pellets were then embedded in fresh Epon in a mold at 60 °C in an oven over 48 h. Ultrathin sections of 200 nm thickness were obtained using a UC6 ultramicrotome (Leica) and mounted on Si wafer supports (Ted Pella, USA).

All datasets were produced by imaging samples with an Axia ChemiSEM (Thermo Fisher Scientific, USA) using a concentric backscattered electron detector (CBS) in beam deceleration mode with a stage bias of −4 kV and landing energies between 2 kV and 4 kV.

The generalization performance of the fine-tuned Cellpose-SAM model for cell segmentation, trained as described in Section 2.4.1, was evaluated qualitatively by comparing its predictions on the 2D SEM datasets with those of the pre-trained baseline Cellpose-SAM model. Predictions were run on images downsampled by integer factors until their pixel width was close to 130 nm. Since the models were operating in 2D mode, no custom flow or mask postprocessing was applied.

#### 2.5.1 Segmentation of nanoparticles

Segmentation of nanoparticles was carried out with a non-parametric Empirical Bayes (EB) segmentation on the full resolution images. The complete data likelihood was modeled as *p* (*I, Z*| *θ*), where *I* the voxel intensity, *Z* ∈ {bg, fg} is class assignment (background vs foreground/nanoparticles), and the *θ* are the model parameters. The empirical intensity distribution *p*_stack_ was obtained by normalizing the stack histogram *h*_stack_, and modeled as a two-component mixture approximating the marginal distribution *p* (*I* |*θ*):

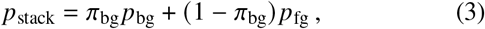

where *p*_bg_ = *p* (*I* |*Z* = bg, *θ*), and *p*_fg_ = *p* (*I* |*Z* = fg, *θ*), are the conditional probabilities of background and foreground, and *π*_bg_ = *p* (*Z* = bg |*θ*) is the background prior. All distributions are non-parametric; the full parameter set is *θ* = {*p*_bg_ [*k*], *p*_np_ [*k*], *π*_bg_} with *k* indexing the histogram bins (intensity levels). Estimation proceeded in two stages: first, *p*_bg_ was estimated via foreground masking (Algorithm 1); then, with *p*_bg_ fixed, *p*_fg_ and *π*_bg_ were estimated via Expectation-Maximization (EM) (Algorithm 2).

The background histogram *h*_bg_ was constructed by per-slice accumulation, excluding foreground regions defined based on a prior. The algorithm is summarized in Algorithm 1. For each slice, an initial foreground mask was created by thresholding at *k*_low_ (99.5th percentile of *p*_stack_). Connected components (CC) smaller than *A*_min_ = 20 pixels were discarded; the bounding boxes (BB) of the remaining blobs were added to the mask with an exclusion margin of *d*_ex_ = 20 pixels that account for the nanoparticles’ point spread function. Finally, all pixels exceeding *k*_high_ (99.8th percentile *p*_stack_) were masked. The remaining background pixels were accumulated into *h*_bg_. Finally, *p* (*I*| *Z* = bg, *θ*) = *p*_bg_ was calculated by normalizing *h*_bg_ and kept fixed throughout the rest of the pipeline.

The remaining parameters *π*_bg_ and *p*_fg_ were then estimated with a non-parametric EM algorithm [50–52], summarized in

##### Algorithm 1

Background histogram construction

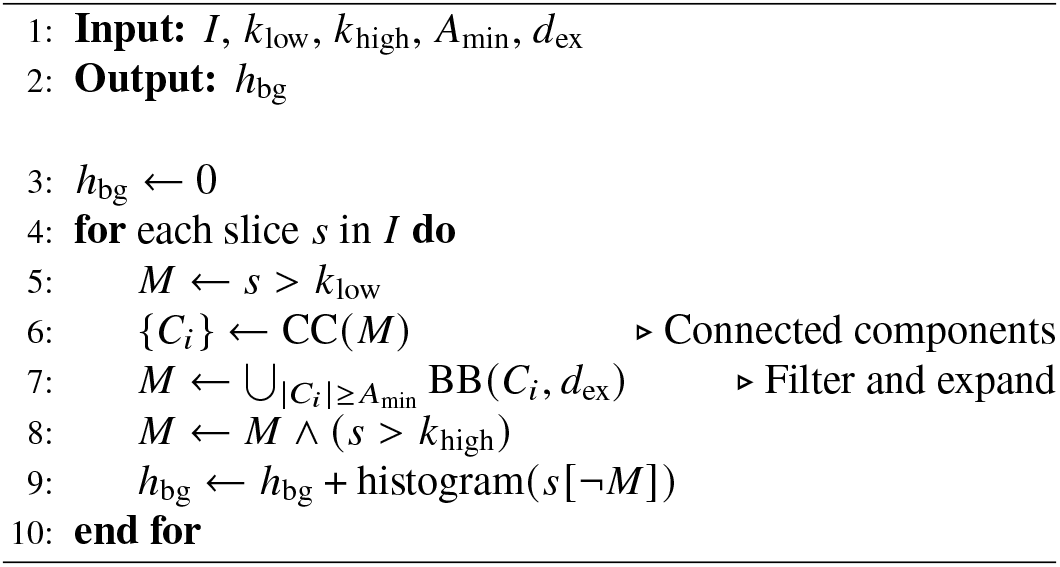

Algorithm 2. The background prior *π*_bg_ was initialized as the ratio of background voxels over total voxels, while the foreground conditional *p*_np_ was initialized as *p*_stack_ above *k*_low_ and uniformly 0 below it. The parameters were then optimized as shown in Algorithm 2. At each iteration, the background responsibility *γ*_bg_ was updated with values from the previous iteration, *p*_fg_ and *π*_bg_ were updated with the newly calculated *γ*_bg_ and the negative log-likelihood (NLL) of the data was calculated. The algorithm was deemed to have converged if the NLL improvement was below a threshold τ = 1 ×10^−10^. All operations are element-wise unless specified.

The foreground posterior probability *p* (*Z* = fg |*I, θ*) = *p*_fg| *I*_ was then calculated by considering each slice as a separate mixture model and applying Bayes’ rule. Equation 4 shows the model for the empirical intensity distribution of each slice *p*_slice_:

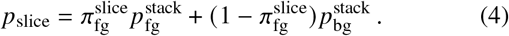

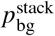 and 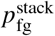are the conditional distributions for foreground and background obtained in the previous steps, with superscripts indicating that they were calculated on the full stack. The slice-specific empirical foreground prior 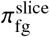 was calculated by maximum a posterior (MAP) estimation under prior 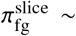 Beta (*α* = 1, *β* = 20) by minimizing the log posterior. The foreground posterior was then calculated as:

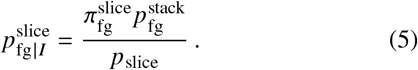

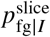 was computed for each intensity bin and applied as a lookup table to obtain a per-voxel posterior map. The operation was repeated for each slice, and the posterior probabilities of the entire stack were smoothed with a non-isotropic Gaussian kernel in logit space and converted back. This step acted as a spatial regularization, and operating on logits ensures smoothed values remain valid probabilities and avoids boundary compression near 0 and 1. The smoothed probability map was then downsampled to isotropic (50 nm× 50 nm× 50 nm) spacing by average pooling.

Nanoparticle masks were finally computed by thresholding the foreground posterior at 0.95. The masks were post-processed by binary closing with a ball element of radius 1, followed by labeling with a connected component analysis (2-connectivity). Labeled objects smaller than 10 voxels were discarded. The number of NPs per detected cluster was estimated by modeling them as equal spheres with 50 nm diameter and assuming close-packing with density 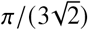; the mass per cluster was then simply obtained by multiplying the number of NPs by their mass *m*_NP_ = 1.26 × 10^−18^ g.

##### Algorithm 2

Non-parametric EM for mixture deconvolution

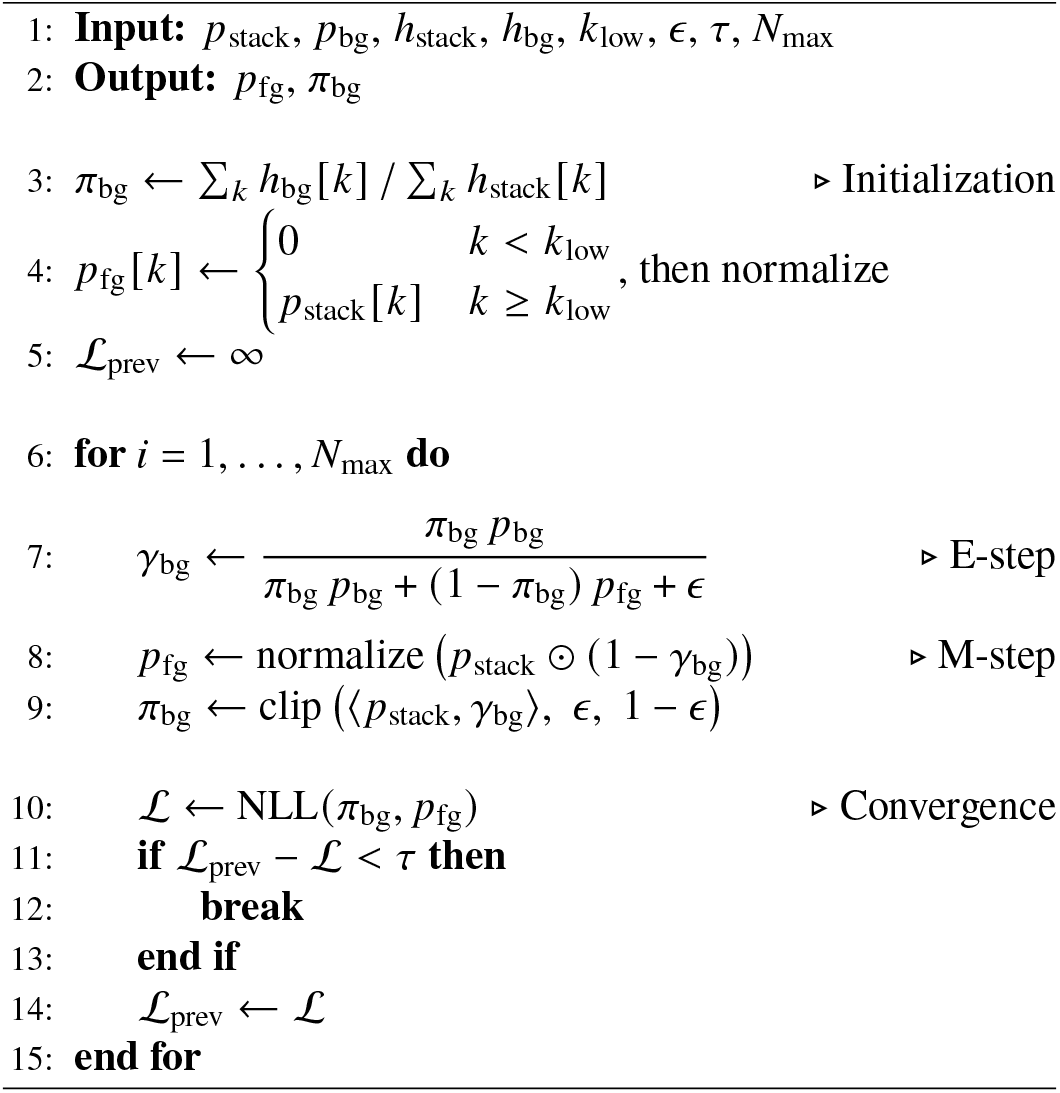

### 2.6 Analysis of cellular structures shape descriptors

Shape descriptors of cellular structures were calculated with a hybrid approach combining voxel-based, mesh-based, and implicit surface representation approaches.

First, each object’s centroid and bounding box were calculated from the labeled voxel image using scikit-image [53] and cuCIM (https://github.com/rapidsai/cucim). The eigenvalues *λ*_1_ ≤ *λ*_2_ ≤ *λ*_3_ of the object’s gyration tensor were calculated as well. The object aspect ratio was then calculated as 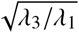

An implicit representation of each object’s surface was then obtained as the 0-level set of the signed distance function (SDF) *ϕ* from the object boundary, with *ϕ* < 0 on the inside [54, 55]. The SDF was calculated from the directly from the object voxel mask, and smoothed with a Gaussian filter with σ = 3 to remove step artifacts arising from original voxel boundary. The object volume was calculated simply as the count of voxels for which *ϕ* < 0. The object surface area was calculated using a delta function approach as [54]:

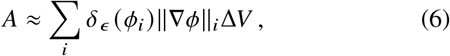

where Δ*V* is the voxel volume, *i* is the voxel index, and δ_ϵ_ is the derivative of the smeared-out Heavyside function:

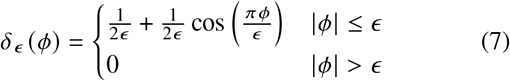

A bandwidth parameter ϵ = 75 nm was used for the computation. Sphericity was calculated as Ψ = *π*^1/3^(6*V*)^2/3^/*A* and equivalent diameter as *d*_eq_ = 2(3*V*/4*π*)^1/3^.

The smoothed SDF was then used to generate a mesh using the marching cubes algorithm at *ϕ* = 0 [56]. The SDF was downsampled by a factor 2 prior to this step to reduce the number of mesh vertices while still maintaining an accurate representation of the surface. The mesh vertices were then used to sample the gradient ∇*ϕ* and Hessian *H*_*ϕ*_ of the full-resolution *ϕ* on the surface at sub-voxel resolution. First and second derivatives were computed using central finite differences with step *h* = 1.5 in voxels. The mean *K*_M_ and Gaussian *K*_G_ curvatures were then calculated as [57]:

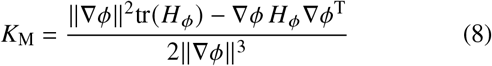

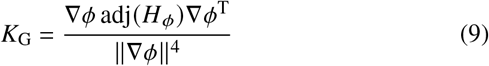

The principal radii of curvature κ_1_, κ_2_, and their derived properties curvedness *C* and shape index *S* [29] were then calculated as:

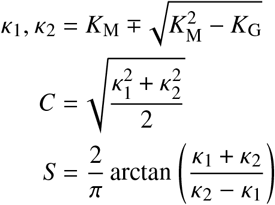

The local measures of curvature *K*_M_, *K*_G_, *C*, and *S* were aggregated per object using the vertex areas as weights, in order to account for the non-uniform density of mesh vertices on the surface. Vertex area was calculated as 1/3 of the area of each mesh face touching the vertex.

The Minkowski-Bouligand fractal dimension *D*_MB_ was calculated using the tube volume approach directly on *ϕ* [58, 59]. For this calculation, *ϕ* was smoothed with a Gaussian filter with a smaller standard deviation than in the previous steps (σ_frac_ = 0.7), in order to balance voxel artifact removal with loss of fine structure. The volume of tubes *V* (*r*) = Σ _|*ϕ*| <*r*_ Δ*V* was calculated for *n* = 30 logarithmically spaced *r* ∈ [*r*_min_, *r*_max_]. The bounds of *r* were defined as *r*_min_ = max(3σ_frac_Δ*x*, 1.5Δ*x*) and *r*_max_ = *d*_eq_/4, where Δ*x* is the voxel pitch. *D*_MB_ is defined as:

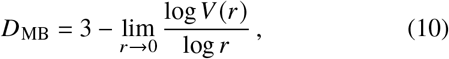

and was calculated from the slope *s* of the linear part of the log-log plot of *V* (*r*) as a function of *r*, as *D*_MB_ = 3 − *s*. To account for finite-size saturation at large *r*, points were iteratively trimmed from the upper end to maximize the *R*^2^ of the linear fit, down to a minimum of 5 points.

### 2.7 Analysis of nanoparticle-nucleus distance

The distance from the nucleus of each cell was calculated by applying an Euclidean distance transform (EDT) to a binary mask of the nuclei contained in the cell. Values outside the cell and values in cells where no nucleus was segmented were masked to NaNs. The distance of each nanoparticle voxel from the closest nucleus in the corresponding cell was then calculated by a simple indexing of the calculated EDT.

### 2.8 Statistical analysis

All statistical tests were two-sided. Mann-Whitney U tests were used to assess the independence of aggregated Jaccard index populations across segmentation models; *t*-tests on interaction and category terms of linear regression models were used to compare regression parameters across object classes. A threshold of *p* < 0.05 was used to determine statistical significance. Statistical analyses were performed using SciPy v1.17.1 [60], statsmodels v0.14.6 [61], and statannotations v0.7.2 [62]. Unless otherwise stated, quantities are reported as mean ± standard deviation or median [Q25–Q75]. Box plots show the median (center line), interquartile range (box), and 1.5 × IQR (whiskers); individual data points are overlaid as a strip plot. All analyses in this study are based on a single imaged spheroid; no biological replication was performed.

## 3 Results and discussion

A conceptual overview of the whole analytical workflow is shown in Figure 1.a: we first cultured Au-spiked FaDu spheroids following a previously published protocol [31]. We used very low cell seeding density to grow small tumor spheroids (225 µm ±15 µm, (n=9)), deliberately chosen so that the SBF-SEM field of view would constitute an appreciable fraction of the total spheroid volume. We selected one spheroid and imaged a volume of approximately 102 µm ×102 µm× 35 µm close to its core (Figure 1.c), corresponding to 6.1 % of the total spheroid volume; for reference the same imaged volume would have constituted only 0.3 % of the total volume of a spheroid with a 400 µm diameter [31]. A qualitative observation reveals that nanoparticles occur primarily in clusters within cells, and that the imaged cells have a wide range of shapes. Moreover, staining was effective in providing contrast to structures of interest (membranes, organelles), and the cellular ultrastructure appears well preserved.

**Figure 1:**
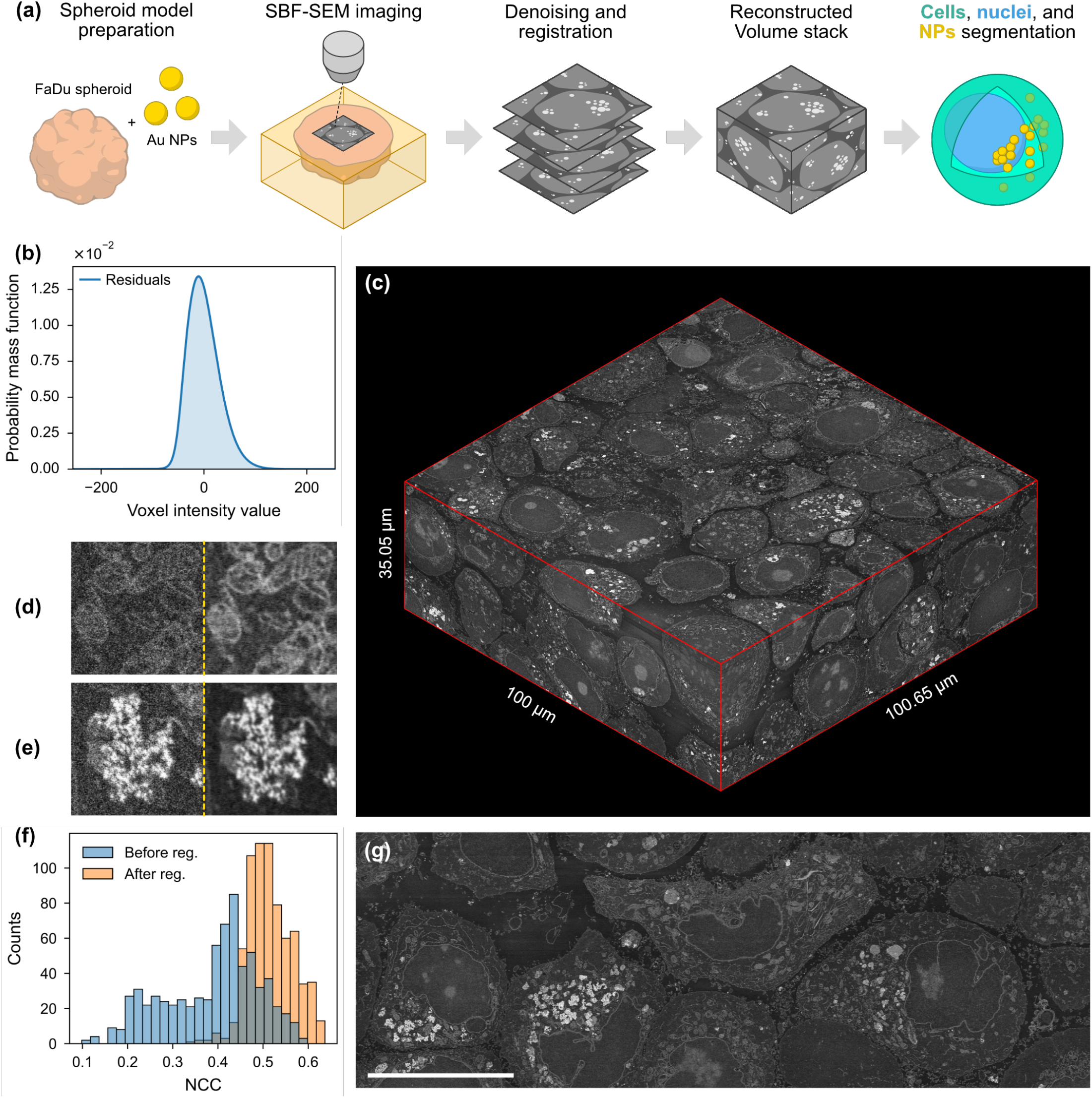
(a): Conceptual overview of the work. NP illustration adapted from BioIcons (bioicons.com, Rahel-Khdhr, CC BY 4.0). (b): Distribution of image residuals, calculated as the difference between the denoised and the noisy image and accumulated over the entire volume. (c): 3D representation of the imaged volume after denoising and registration. The volume was automatically cropped to remove empty borders introduced by image warping, yielding final dimensions of 100.00 µm ×100.65 µm × 35.05 µm. (d-e): Close-up view of organelles (d) and NPs (e) before (left) and after denoising (right). (f): Histogram of normalized cross correlation (NCC) between each neighboring *XY* image pair before and after registration (*n* = 700). (g): Full size *Y Z* section (*x* = 33 µm) of the registered volume, showing the good quality of the alignment; scale bar: 20 µm.

### 3.1 Preprocessing

An appropriate preprocessing pipeline is a key component of a quantitative image analysis workflow, and this is particularly true for datasets acquired with volume EM techniques [8].

We first denoised the images with Noise2Void, a state-of-the-art deep learning algorithm for self-supervised noise removal [34]. Training curves for the model on our data are presented in Figure S1, highlighting a fast learning behavior and no overfitting. Figure 1.b shows the distribution of image residuals *I*_res_ = *I*_noisy_− *I*_denoised_ across the entire volume stack, which is a proxy for the learned noise distribution. The distribution is smooth and roughly centered around 0 with a heavier tail at higher values. While not readily parameterized by simple analytical distributions, this empirical noise model supports a consistent denoising process and reflects the complex signal-dependent noise characteristics of SEM imaging [63]. The quality of the denoising process can also be evaluated by examining high-magnification details of images before and after denoising, as shown in Figure 1.d-e. Noise appears indeed efficiently filtered out, and image features for both cellular structures and nanoparticles are preserved without introducing evident artifacts.

Moreover, while SBF-SEM in principle produces inherently aligned images, small differences in image orientation can still be present, as well as subtle warping of the resin block. If not corrected, this effect can lead to inaccurate object masks during segmentation. This is especially relevant for nanoparticles, since the *Z*-resolution of the dataset (50 nm) is not high enough to resolve individual 50 nm particles, and even small voxel shifts can radically alter the shape of a small aggregate. We therefore registered the 2D images in the volume stack using an intensity-based criterion that maximizes the normalized cross correlation ρ_NCC_ between each neighboring image pair. This approach has a higher computational cost than sparse feature-based methods, which are more commonly used in the field [8]. However, the near-identical image content between neighboring slices and small displacement fields make the case for an intensity-based approach, which has the advantage of not relying on manually-extracted features [64]. To reduce the computational load of our approach, we developed a light GPU implementation that relies on multi-scale image pyramids. A typical optimization curve is shown in Figure S2.a for a single image pair; note that optimization is implemented as a minimization of ℒ_pair_ = 1 −ρ_NCC_. As expected, most of the improvement occurs at the coarser scales, while the more expensive finer ones provide minimal fine adjustments. The ρ_NCC_ distribution is much narrower and shifted to higher values after registration (*µ* = 0.39, *σ* = 0.10 before vs *µ* = 0.51, *σ* = 0.05 after), as shown in Figure 1.f, which demonstrates quantitatively that there is a clear improvement in image alignment. Moreover, it can be verified that this improvement occurs for every image pair (Figure S2.b). This can also be visualized directly on the registered stack, as shown in the *X Z* and *Y Z* slices of the 3D volume render (Figure 1.c) and in selected full-size *Y Z* cross section (Figure 1.g). No major misalignment can be detected upon visual inspection, which further demonstrates the effectiveness of our approach. After registration, the exact *XY Z* size of the volume is 100.00 µm × 100.65 µm × 35.05 µm.

### 3.2 Segmentation of cells, nuclei, and nanoparticles

The spheroid dataset has two main families of objects to segment: cellular structures (cells, nuclei) and nanoparticles. This presents a unique challenge: while cellular structures have a typical equivalent diameter of a few µm and possess a wide array of features, nanoparticles in this experiment have a diameter of 50 nm (although they usually occur in larger clusters) and are primarily identified by their voxel intensity. Moreover, a thorough labeling of nanoparticle ground truth voxels would be unfeasible due to their small size and sheer number. Based on these considerations, we followed a hybrid approach where each target is segmented following a different strategy:

- Cellular structures were segmented by fine-tuning a pre-trained deep learning model (Cellpose-SAM [15]) on a small labeled subset of our data. Separate models were trained for cells and nuclei.
- Nanoparticles were segmented using a non-parametric empirical Bayes approach, which avoids the need for labeled training data or complex deep learning architectures.

Training independent models also allowed for a straightforward overlapping class assignment, which is necessary for our analysis.

#### 3.2.1 Segmentation of cells and nuclei

We first evaluated our segmentation approach by comparing the accuracy of Cellpose-SAM, fine-tuned to our data, against the baseline pre-trained model (Figure 2.c-d, summarized in Table 2).

**Table 2:** Segmentation metrics by target and model. Aggregated Jaccard Index (AJI) is reported as mean ± SD.

| Target | Model | AJI | AP$_{\tau=0.5}$ |
| --- | --- | --- | --- |
| Cells | Fine-tuned | $0.940 \pm 0.020$ | 0.830 |
| | Pre-trained | $0.670 \pm 0.115$ | 0.499 |
| Nuclei | Fine-tuned | $0.936 \pm 0.043$ | 0.864 |
| | Pre-trained | $0.272 \pm 0.062$ | 0.100 |

**Figure 2:**
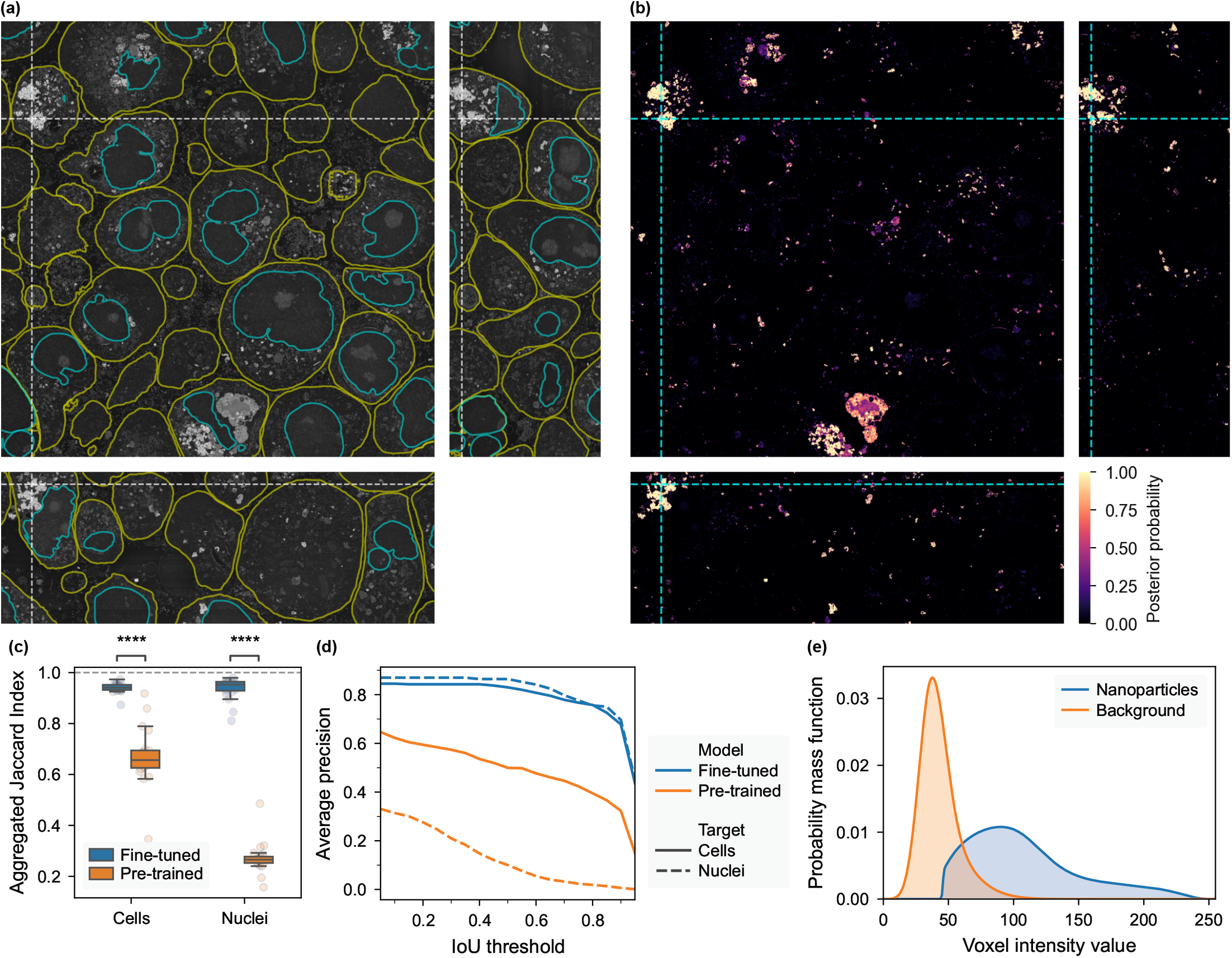
Segmentation of cellular structures and nanoparticles. (a) Orthogonal cross-sections overlaid with cell (yellow) and nucleus (cyan) outlines. (b) Orthogonal cross-sections of the nanoparticle posterior probability maps. (c–d) Aggregated Jaccard Index (AJI) (c) and average precision (AP) at varying intersection over union (IoU) thresholds (d) comparing the fine-tuned and pre-trained Cellpose-SAM models for cell and nucleus segmentation, evaluated on the full labeled dataset (*n* = 20 images); ****: *p* ≤ 10^−4^, Mann–Whitney U test. (e) Voxel intensity probability mass functions for the nanoparticle and background components estimated by expectation–maximization (EM) decomposition.

The fine-tuned model substantially outperformed the pre-trained baseline for cell segmentation, both in terms of average precision (representing instance detection accuracy) and aggregated Jac-card index (indicating voxel-level instance delineation accuracy) relative to the labeled ground truths. The benefits of fine-tuning were even more pronounced for nuclei segmentation, though the baseline model was not originally trained for this task. These results are particularly notable given that Cellpose-SAM was trained on light microscopy data, demonstrating strong generalization to electron microscopy with limited fine-tuning. When outlines of the segmented cells and nuclei were superimposed to the original image, they showed excellent visual agreement with the raw data (Figure 2.a). Our model also outperformed the benchmark experiment performed with Amira at all AP threshold values (Figure S4, Table S1). For cell segmentation, it further yielded a small but significant improvement in the AJI (*p* = 4.6× 10^−4^), whereas the improvement for nuclei, while present, was not significant (*p* = 0.057).

In general, an important limitation of Cellpose-SAM is that the objects to segment should have a characteristic size comparable to that of the tiles used by the model for prediction (256 × 256). For our dataset, this means that the model cannot be run at the full *XY* resolution, where cells can have diameters in the order of thousands of pixels, but must be applied to a downscaled version of the data and the predicted labels need to be upscaled. Despite this, our approach achieved remarkable results at the resolution used for the analysis.

We tested the generalization of our fine-tuned Cellpose-SAM model qualitatively by comparing its predictions against the pre-trained baseline model on a variety of 2D SEM datasets, overlaid on the original images (Figure S5, Table 1). The fine-tuned model produced generally more accurate segmentation for all tested cases, although the extent of this improvement varied greatly across datasets. For the HT1080-Hf and FaDu-Au datasets, the baseline model achieved already very good results, and fine-tuning mainly provided improved contour accuracy and fewer spurious detections. A549-UCNP is the dataset where improvement due to fine-tuning was the most evident: the baseline model had several false negatives and partial detections, while the fine-tuned one correctly segmented all cells. On the other hand, for the MT-Ti-MIL dataset, our model showed its limits: while the fine-tuned model still produced better results than the baseline, the overall segmentation quality was quite poor. The lack of generalization to this dataset can be readily explained as domain shift, as the dataset features and contrast were substantially different to those of the training dataset. Nevertheless, the good performance on the other 2D datasets as well as the results of fine-tuning on the vEM data supports the hypothesis that, with additional training on a larger and more varied dataset, Cellpose-SAM could be adapted to be a generalist cell segmentation model for SEM data.

#### 3.2.2 NP segmentation

Following Equation 3, we decomposed the empirical intensity distribution of the entire volume stack *p*_stack_ into the calculated background *p*_bg_ and foreground (nanoparticle) *p*_fg_ distribution, as shown in Figure 2.e. The non-parametric EM algorithm (Algorithm 2) we used for the decomposition converged to *w*_bg_ = 0.982 with ℒ= 4.103. This decomposition allowed us to calculate a separate posterior probability for each slice in the stack, which accounts for variations in foreground density and slight illumination shifts. We then used this to build a nanoparticle probability map, where every voxel is assigned a value representing the probability that it belongs to a nanoparticle, as shown in Figure 2.b. Brighter areas are correctly assigned high nanoparticle probability, while intermediate features (e.g., organelles or stain precipitates) receive lower values, and background approaches zero. Since, according to our method, probability assignment is purely based on intensity, we chose a very high confidence threshold (95 %) for the final nanoparticle segmentation to remove spurious detections from other bright objects. Figure S6 shows a high-magnification thresholded map, illustrating good agreement between nanoparticle aggregates and segmented regions. Isolated nanoparticles and very small aggregates were excluded, as the *Z*-resolution is insufficient to resolve them and they cannot be reliably distinguished from noise.

### 3.3 Analysis of segmented masks

#### 3.3.1 NP distribution

Combining the segmented masks of cells, nuclei, and nanoparticles allowed us to fully reconstruct their relative position and investigate their spatial relationships. For the purpose of studying cell-nanomaterial interactions, representing these data on a per-cell basis is a natural choice, and a few examples of this are shown in Figure 3.a-c, where individual nanoparticles clusters, as identified by connected component analysis, are labeled in different colors. Most clusters were found to be comparatively small, with a median [Q25–Q75] estimated NP count of 76 [27– 385]; however, a few clusters were extremely large, estimated to contain up to millions of NPs, and appeared as complex inter-connected 3D networks (Figure 3.c). It should be acknowledged that inaccuracies in the connected component labeling might bias these results by merging neighboring clusters, but a visual inspection of the original images supports our observations. It has been suggested that similar micron-sized structures arise from the conglomeration and fusion of endocytotic vesicles in the perinuclear region, in the vicinity of the endosomal recycling compartment [65, 66]. Future studies combining vEM segmentation with trafficking markers could leverage this approach to investigate the underlying mechanisms.

**Figure 3:**
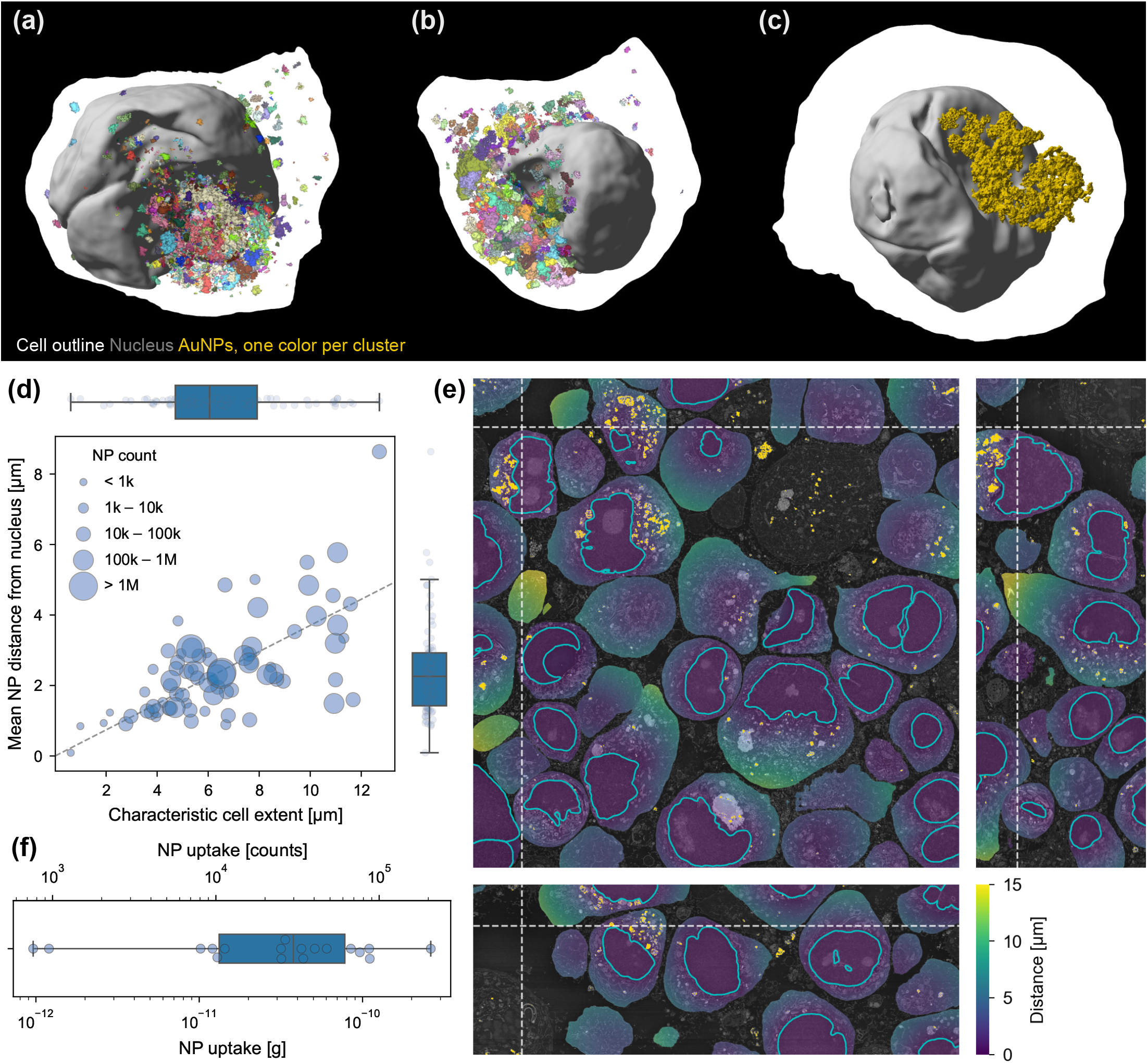
3D evaluation of nanoparticle uptake and distribution. (a-c): 3D representations of selected cells with their nucleus and internalized NPs, shown to scale; each NP cluster is rendered in a distinct color. In (c), only a single cluster is displayed to highlight the complex morphology of large, micron-sized clusters. (d): Scatter plot of mean NP distance from the host cell nucleus as a function of characteristic cell extent (*n* = 78); marker size encodes the estimated NP count internalized per cell; dashed line: linear regression (slope 0.371, intercept 0, *R*^2^ = 0.404). (e): Orthogonal cross-sections of the imaged volume, superimposed with a map of the distance from the nucleus for each cell; nuclei are outlined in cyan, and NPs are labeled in gold. (f): Estimated NP uptake by mass and count for cells fully contained within the imaged volume (*n* = 18).

We then characterized the distance of NPs from the nucleus of the cell in which they are internalized (Figure 3.e); cells without a detected nucleus were excluded from the analysis (*n* = 78). NPs appeared primarily concentrated close to the nucleus, with a median distance of 2.57 µm [1.17 µm–3.78 µm] (Figure S7.c) and a mode at approximately 0.8 µm; no nanoparticles were detected inside the nucleus. To study whether NP position within the cell depended on cell size, we aggregated the distance of NPs from the nucleus per cell and we plotted it against a characteristic cell extent, (Figure 3.d), which we calculated as the 95^th^ percentile of the distance from the nucleus computed over all voxels within the cell; furthermore, we encoded information about the estimated count of internalized NPs via the plot marker size. We used the characteristic cell extent as a proxy for cell size because most cells were not fully inside the imaged volume, causing straightforward parameters such as cell volume or equivalent diameter to be systematically underestimated. We found a moderate correlation between the two variables (*R*^2^ = 0.404), which suggests that larger cells, having more space to accommodate NP clusters, might have a wider distribution of NP positions. This is also directly supported by evaluating the spread of NP distance from the nucleus as a function of characteristic cell extent (Figure S7.a), which shows a moderate positive correlation as well (*R*^2^ = 0.568). Nevertheless, for most cells, NP clusters still tend to be primarily concentrated relatively close to the nucleus, regardless of the estimated internalized NP count (Figure 3.d and Figure S7.b). The agreement between the two median values (2.26 µm per cell vs 2.57 µm globally) suggests that high-uptake cells, despite containing orders of magnitude more NPs, do not show a systematically different spatial distribution. Similar clustering behavior in the perinuclear region has also been reported in the literature. For example, HfO_2_ nanoparticle agglomerates in Caco-2 cells showed pronounced perinuclear accumulation, with approximately 50 % located within 1 µm and approximately 75 % within 2 µm of the nuclear membrane [67], whereas SPION-containing vesicles in MCF-7 breast cancer cells were reported at nanoparticle-to-nucleus distances ranging from 2.3 to 3.8 µm [66]. The quantitative differences across systems likely reflect the influence of nanoparticle formulation, surface chemistry, aggregation state, and cell type on subcellular localization, even when the overall perinuclear accumulation pattern is preserved. These differences in intracellular distribution may contribute to differences in therapeutic efficacy and are important for understanding the nanoparticles’ mode of action: for example, in NP-mediated radioenhancement, enhanced cell damage (secondary electron emission and reactive oxygen species generation) is highly local and dependent on the relative distance of NPs to a biological target, such as DNA [68].

From the estimated count of internalized NPs, discussed in the previous paragraph, we estimated the Au mass uptake for each cell (Figure 3.f). We limited this analysis only to cells that were fully visible in the imaged volume (*n* = 18), and we obtained a median NP uptake of 0.042 ng/cell [0.013 ng /cell–0.085 ng /cell]. Moreover, despite the small number of cells considered, we observed that the estimated uptake values span several orders of magnitude, which is reflective of the well-known heterogeneity in NP uptake by cells [27, 28]. Inductively Coupled Plasma Optical Emission Spectroscopy (ICP-OES) data of similar experiments [31] report uptake of roughly 0.05 ng /cell for FaDu cells exposed as a monolayer but otherwise similar conditions (50 nm Au NPs, 50 µg mL^−1^ exposure concentration), which corresponds to our initial exposure prior to spheroid formation. While bulk ICP-based techniques enable sensitive quantification of total elemental uptake, they do not resolve intracellular heterogeneity, distinguish between particulate and dissolved species, or determine proximity to relevant biological targets such as the nucleus. Thus, while vEM and ICP should be regarded as complementary approaches, vEM is particularly informative in cases where therapeutic effects are governed by local subcellular interactions rather than total elemental content.

#### 3.3.2 Cellular shape descriptors

In order to analyze the shape of cells and nuclei in detail, we calculated high-resolution triangle meshes using the marching cubes algorithm. Representative meshes are shown in Figure 4.a-f using the calculated local mean curvature as texture.

**Figure 4:**
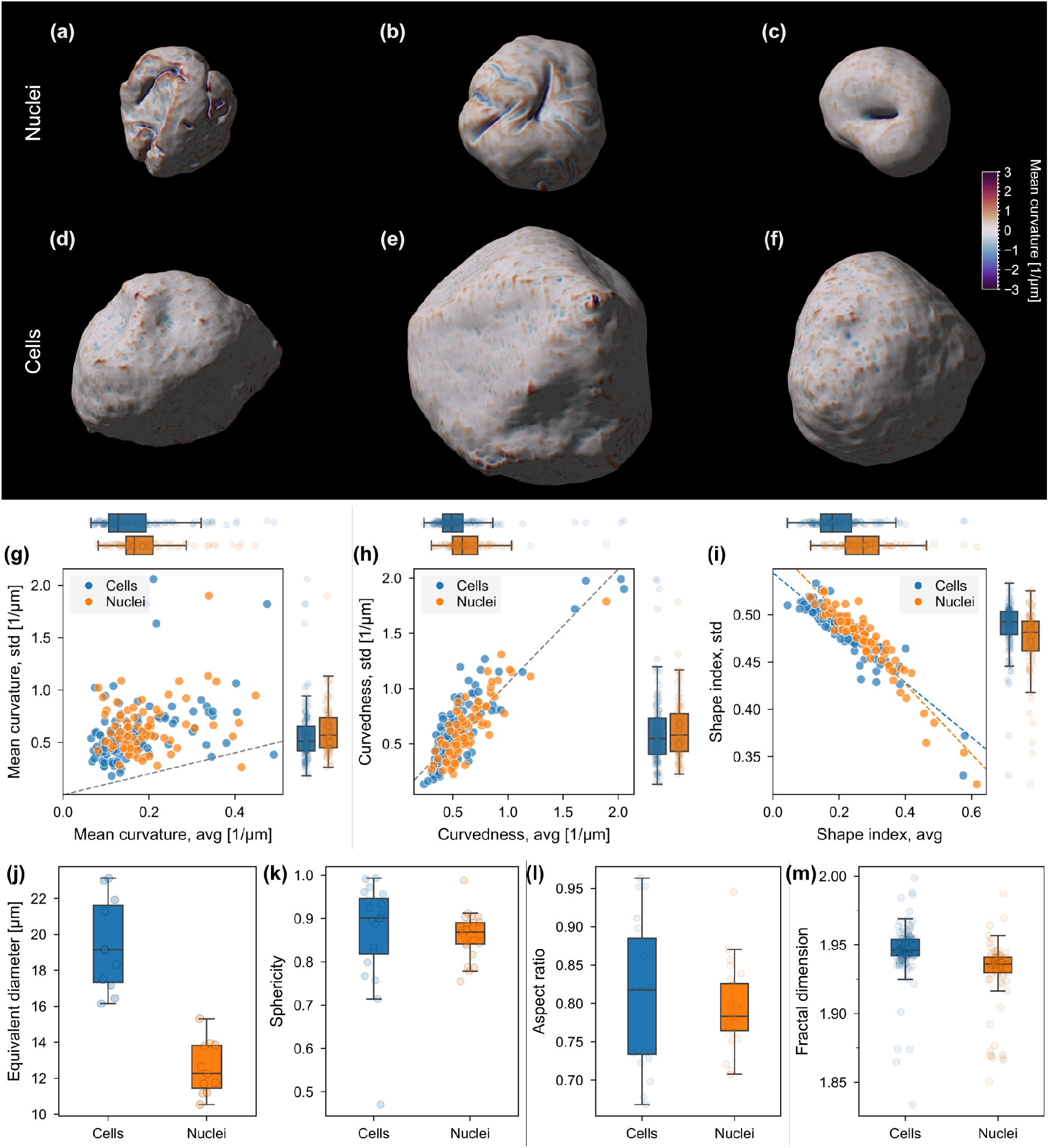
Evaluation of 3D cell and nuclei shape and curvature descriptors. (a-f): 3D renderings of selected cells (d-f) and their corresponding nuclei (a-c) with mean curvature mapped on their surface, to scale. (g-i): Scatter plots of standard deviation (std) as a function of the average value (avg) of mean curvature (g), curvedness (h), and shape index (i), aggregated per cell (*n* = 124) and nucleus (*n* = 83); dashed lines: (g) reference line at *x* = *y*; (h): linear regression (slope 1.03, intercept 0.019, *R*^2^ = 0.738); (i): linear regressions for cells (blue, slope −0.29, intercept 0.54, *R*^2^ = 0.839) and nuclei (orange, slope −0.37, intercept 0.57, *R*^2^ = 0.894). (j-l): Box plots of global shape descriptors for cells and nuclei fully contained within the imaged volume: (j): equivalent diameter (*n* = 11 cell–nucleus pairs); (k): sphericity (*n* = 20 cells, *n* = 24 nuclei); (l): aspect ratio (*n* = 20 cells, *n* = 24 nuclei). (m): Estimated fractal dimension for all cells (*n* = 124) and nuclei (*n* = 83).

We averaged the mean curvature for each cell (*n* = 124) and nucleus (*n* = 83) weighted by mesh vertex area and report it alongside its standard deviation to capture the spread of local curvature (Figure 4.g). The obtained values of the averaged mean curvature are of difficult interpretation due to the signed nature of the parameter [69]: grooves and ridges cancel out, and most cells and nuclei are dominated by large low-curvature regions. For our data, cells and nuclei yielded very similar results, showing positive average mean curvatures, indicating a slight dominance of convex regions. Standard deviations usually exceeded the average, which further highlights their complex shape. In order to decouple curvature shape and magnitude, we also calculated curvedness *C* and shape index *S* from the principal radii of curvature [69]. *C* represents the magnitude of local curvature (Figure 4.h), and we observed a moderately strong positive linear correlation between its average and standard deviation values for our data (*R*^2^ = 0.738). This can be interpreted as objects in our dataset having more pronounced curvature also showing higher variability, regardless of their local shape type. Moreover, neither the slope nor the intercept differed significantly between cells and nuclei, as determined by *t*-tests on the interaction and category terms of a linear regression model (*p* = 0.732 and *p* = 0.193, respectively), justifying a single pooled regression.

Analyzing *S*, which represents the local shape type of the surface, with 1 being spherical cap and −1 being spherical cup, yields additional insight (Figure 4.i). We observed a strong negative linear correlation between *S* average and standard deviation; moreover, cells and nuclei followed two slightly different linear trends (*R*^2^ = 0.839 and *R*^2^ = 0.894, respectively), and the difference in their intercept and slope is statistically significant (*p* < 0.05 for both, determined as above). We interpreted these results as cells and nuclei with a more regular shape having more locally spherical-cap-like character (low standard deviation, high average *S*), while more complex shapes are associated with higher variability. Furthermore, the slightly steeper slope of the nuclei trend compared to cells (−0.37 *vs* −0.29) suggests that the relationship between surface regularity and local shape type is more pronounced in nuclei than in cells. This is also consistent with visual inspection, as shown in Figure 4.a-f, where nuclei with intricate shapes typically show complex ridges and invaginations (Figure 4.a-c), features generally absent from cells (Figure 4.d-f).

In addition to studying local curvature, we also calculated global shape descriptors for cells and nuclei. For most of these parameters (equivalent diameter, sphericity, aspect ratio), it is necessary to have a representation of the entire object, and we therefore limited the analysis only to cells and nuclei that were fully inside the imaged volume. It should be kept in mind that, while this prevents our results from being polluted with incorrect estimations, it might introduce an implicit selection bias towards smaller objects, as larger ones have a higher chance of being cut by the imaging boundaries. We first compared the equivalent diameter of cells and nuclei (Figure 4.j), obtaining an average of 19.4 µm for cells and 12.6 µm for nuclei. For equivalent diameter calculation, we further restricted the analysis to cells that were fully visible in the volume and whose nucleus was detected (*n* = 11). We then calculated the sphericity (Figure 4.k) and aspect ratio (Figure 4.l) of cells and nuclei, which describe how close their shape is to a sphere in terms of their surface to volume ratio and principal axes of inertia, respectively. Both parameters yield values close to unity for both cells and nuclei, indicating shapes broadly consistent with a sphere. However, cells show a much higher variability according to both measures. Finally, we calculated the Minkowski-Bouligand fractal dimension *D*_MB_ of the cellular objects (Figure 4.m). Since this calculation does not require the entire surface, we computed this for all the objects, even those that were not fully inside the imaged volume. The estimated fractal dimension was below 2 for all objects, indicating that neither cells nor nuclei exhibit fractal complexity at the resolution of our analysis.

## 4 Conclusions

In this work, we addressed the lack of transparent and reproducible quantitative analysis pipelines for volumetric EM data of cells containing NMs by developing an end-to-end workflow applied to tumor spheroids spiked with AuNPs, and releasing the underlying annotated dataset as a public resource. The dataset covers a volume of 100 ×100 × 35 µm^3^ at 10 ×10 ×50 nm^3^ acquisition resolution, comprising 124 cells and 83 nuclei, and constitutes a useful benchmark resource for the community. Careful sample preparation and image preprocessing, including denoising and stack registration, produced a well-aligned, high-quality image stack suitable for downstream analysis of cells, nuclei, and NP features. Segmentation followed a hybrid strategy: a fine-tuned Cellpose-SAM model for cells and nuclei, and an empirical Bayes approach for NPs. Cell and nucleus segmentation achieved improved accuracy over both the baseline pre-trained model and the AI-assisted segmentation tool in Amira, as assessed by average precision and aggregated Jaccard index. Since Amira relies on a human-in-the-loop workflow, the benchmark reflects one specific segmentation outcome rather than an upper bound on what the approach can achieve. The fine-tuned model generalized well to 2D EM datasets of similar appearance, but performance degraded on data with substantially different image characteristics, highlighting the expected dependency on training distribution. These results suggest that, with a more diverse training set, Cellpose-SAM could be used as a general-purpose segmentation model for EM images of biological samples. Simultaneous segmentation of all three targets enabled comprehensive quantitative characterization of NP distribution and cellular morphology. NPs were found predominantly in the perinuclear region, with a median distance of 2.57 µm [1.17 µm–3.78 µm] from the nucleus, while estimated Au mass uptake spanned several orders of magnitude across cells, consistent with the well-documented heterogeneity in NM internalization. Morphological analysis of cells and nuclei revealed broadly spherical but geometrically complex shapes, with 3D shape descriptors and local curvature metrics providing quantitative access to features that are inaccessible from single sections. Together, these results provide a fully documented, reproducible framework for the joint analysis of NM distribution and cellular morphology in volumetric EM data, with broad applicability to quantitative studies in nanomedicine.

## Acknowledgments

We acknowledge the Scientific Center for Optical and Electron Microscopy (ScopeM) of the ETH Zurich as well as the Center for Microscopy and Image Analysis (ZMB) of the University of Zurich for providing access to their microscopes and analytical software. We wish to thank Dr. Andres Käch for the support throughout the acquisition of the vEM data, Lukas Häuser for revising the mathematical formalism of the methods section, and Dr. Elina Andresen for supplying the NaYF_4_:Er,Yb nanoparticles. We also wish to thank Stephanie Eitner for the support during cell culture and sample preparation. This work was funded by the project MetrINo (23.00360, 22HLT04), which has received funding from the European Partnership on Metrology, co-financed by the European Union’s Horizon Europe Research and Innovation Programme and SERI (REF1131-52104). We acknowledge the Swiss National Science Foundation (Eccellenza grant no. 181290).

## Author contributions

DB: Investigation, Methodology, Formal analysis, Software, Visualization, Data curation, Writing – Original Draft; LRHG: Investigation, Supervision, Writing – Review & Editing; SH: Investigation, Writing – Review & Editing; JMMM: Investigation, Writing – Review & Editing; MSL: Investigation, Writing – Review & Editing; JR: Investigation, Writing – Review & Editing; MF: Resources, Writing – Review & Editing; URG: Resources; VMK: Investigation, Supervision, Writing – Review & Editing; MR: Funding acquisition, Writing – Review & Editing; AG: Conceptualization, Supervision, Funding acquisition, Writing - Original Draft, Writing – Review & Editing; IKH: Conceptualization, Supervision, Funding acquisition, Resources, Writing – Review & Editing

## Data availability

The spheroid dataset together with cell, nucleus, and nanoparticle segmentation masks is available at the BioImage Archive (https://doi.org/10.6019/S-BIAD3263), together with the manually labeled ground truth annotations. The code used for the analysis is available at https://github.com/dv-bt/sphero-vem. Model weights for the fine-tuned Cellpose-SAM models are available at Zenodo (https://doi.org/10.5281/zenodo.19616546). Additional data supporting this study are available from the corresponding author upon reasonable request.

## Supplementary Information

### S1 Supplementary Methods

#### S1.1 En bloc staining and resin embedding

Fixed specimens (2.5 % glutaraldehyde in 0.1 mol L^−1^ sodium cacodylate buffer, pH 7.4, for at least several hours) were processed over three days as follows.

##### Day 1

Samples were rinsed three times in 0.1 mol L^−1^ sodium cacodylate buffer (pH 7.4, 10 min each) and transferred to freshly prepared reduced osmium solution (0.075 g K_4_[Fe(CN)_6_] dissolved in 2.5 mL of 0.2 mol L^−1^ cacodylate buffer, then mixed with 2.5 mL of 4 % aqueous OsO_4_) for 60 min on ice. After five rinses in distilled water (3 min each, room temperature), samples were immersed in freshly prepared thiocarbohydrazide solution (1 % w/v in double-distilled water; dissolved at 40 °C for 1 h with agitation every 10 min, then filtered through a 0.22 µm membrane) for 40 min at room temperature. Samples were rinsed again five times in distilled water (3 min each) and incubated in 2 % aqueous OsO_4_ for 90 min at room temperature, followed by five further distilled water rinses (3 min each). En bloc uranyl acetate staining was carried out in 1 % aqueous uranyl acetate overnight at 4 ^°^C.

##### Day 2

Samples were rinsed five times in distilled water (3 min each, room temperature), then three times in distilled water at 50 °C (10 min each). Lead aspartate staining was performed at 50 °C for 2 h using a freshly prepared solution (4 mg mL^−1^ aspartic acid and 6.6 mg mL^−1^ lead nitrate (Pb(NO_3_)_2_) in double-distilled water, adjusted to pH 5.0). Samples were rinsed three times in distilled water at 50 °C (10 min each) and allowed to cool to room temperature. Dehydration was carried out through a graded ethanol series (70 %, 80 %, 96 % and 100 %, 20 min each), followed by two additional incubations in anhydrous ethanol (15 min each) and two incubations in propylene oxide (15 min each). Infiltration was initiated in a 1:2 (v/v) mixture of Epon–Araldite resin and propylene oxide overnight at room temperature.

##### Day 3

Samples were transferred to 100 % Epon–Araldite resin for approximately 50 min and polymerized in embedding molds at 60 °C for 28 h.

#### S1.2 Amira benchmark

The fine-tuned Cellpose-SAM models for cells and nuclei were benchmarked against segmentation results obtained using the commercially available Amira 3D software suite (version 2025.1.1; Thermo Fisher Scientific; RRID: SCR_007353). The benchmark experiment was intentionally designed to balance effort and rigor, requiring approximately 10 hours to generate the ground truth masks, including careful proofreading and refinement of the training data.

The denoised and aligned images, resampled to (50 nm × 50 nm × 50 nm) spacing, were filtered using a 3 × 3 square median filter for one iteration, processing the images slice by slice (*Interpretation*: *XY* Planes, *Iterations*: 1; *Type*: Iterative), before separate models were trained for cells and nuclei.

For the nuclei model, an initial ground truth training mask was generated by segmenting 55 nuclei in three randomly chosen slices of the image stack using the AI-assisted selection tool. Manual refinements were applied where necessary. Nuclei cut off at the edges of the images were also included. Nine sub-regions, each containing at least part of the labeled nuclei, were defined for training the model. The model was trained for 15 epochs using a 2D deep learning network based on the UNet architecture with a VGG19 backbone. Based on the segmentation obtained results, refinements were made by adding more training nuclei to the ground truth mask, mainly in areas where detection was weak. This resulted in a final training dataset of 83 nuclei across 21 sub-regions. The model was then retrained for 15 epochs using the VGG19 network. To distinguish individual nuclei, the segmentation result was represented as instance masks using Amira’s *Labeling* module (*Interpretation*: 3D; *Neighborhood*: 6), with each mask corresponding to a single nucleus.

The cells model was again trained in a similar two-step fashion, starting with an initial ground truth mask containing 41 cells, including those cut off at the edges of the image, in a single slice of the image stack. The model was then trained for 100 epochs using seven partially overlapping sub-regions and the same VGG19 network described above. The ground truth mask was then further adapted to include a total of 109 cells across three image slices. Seventeen partially overlapping sub-regions were used for training, and the model was retrained for 100 epochs. In this case, the *Separate Objects* module was applied to generate an instance mask. This module combines watershed, distance transform, and numerical reconstruction algorithms (*Method*: Chamfer (conservative); *Interpretation*: 3D; *Neighborhood*: 26; *Marker Extent*: 24; *Output Type*: Connected Object; *Algorithm Mode*: Repeatable).

### S2 Supplementary Figures

#### S2.1 Preprocessing

**Figure S1:**
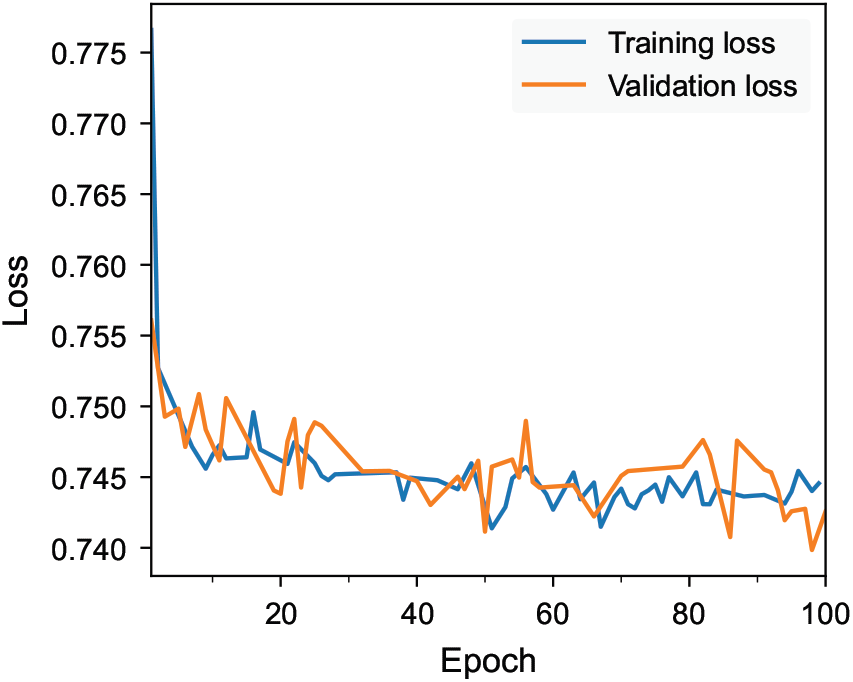
Training and validation losses as a function of epoch of the employed Noise2Void model.

**Figure S2:**
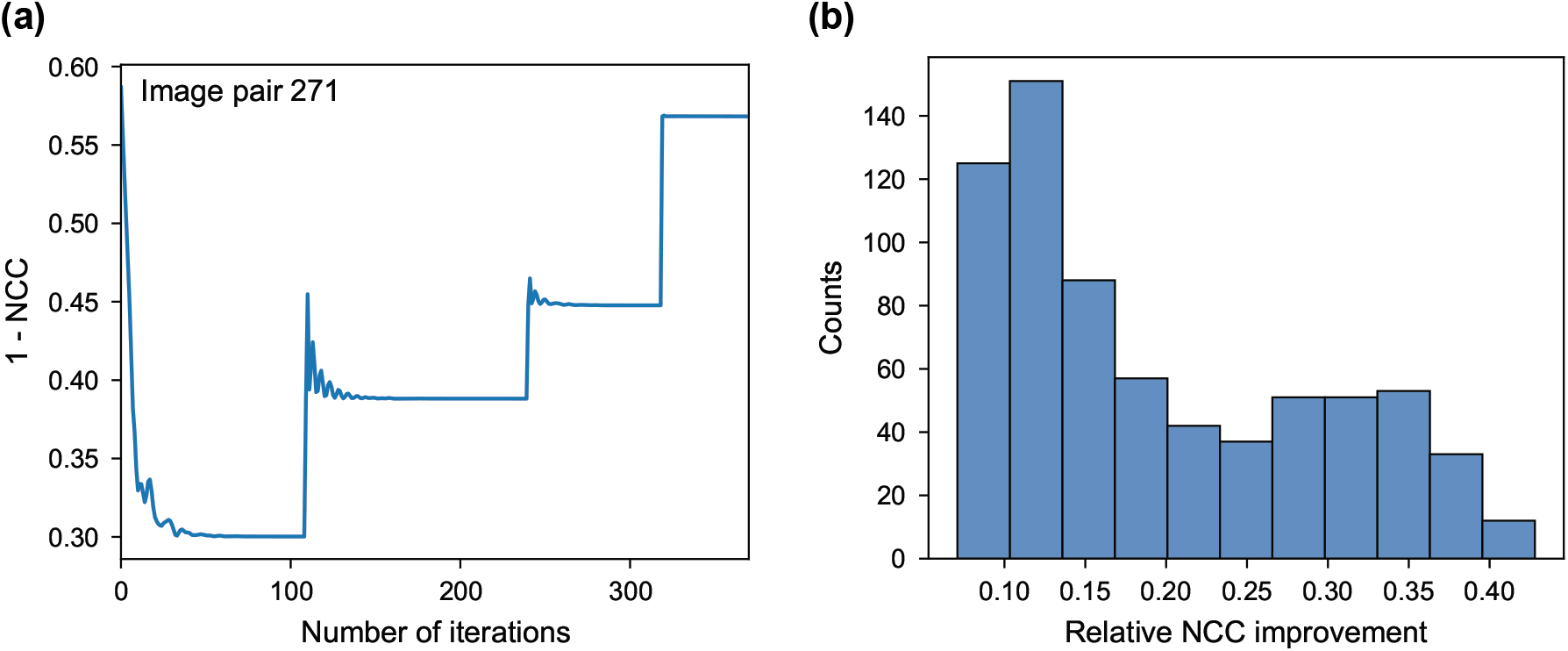
(a): Optimization loss as a function of iteration for the multi-scale registration of one image pair (pair index 271). (b): Relative improvement in normalized cross correlation for each image pair in the volume stack (*n* = 700).

#### S2.2 Segmentation

**Figure S3:**
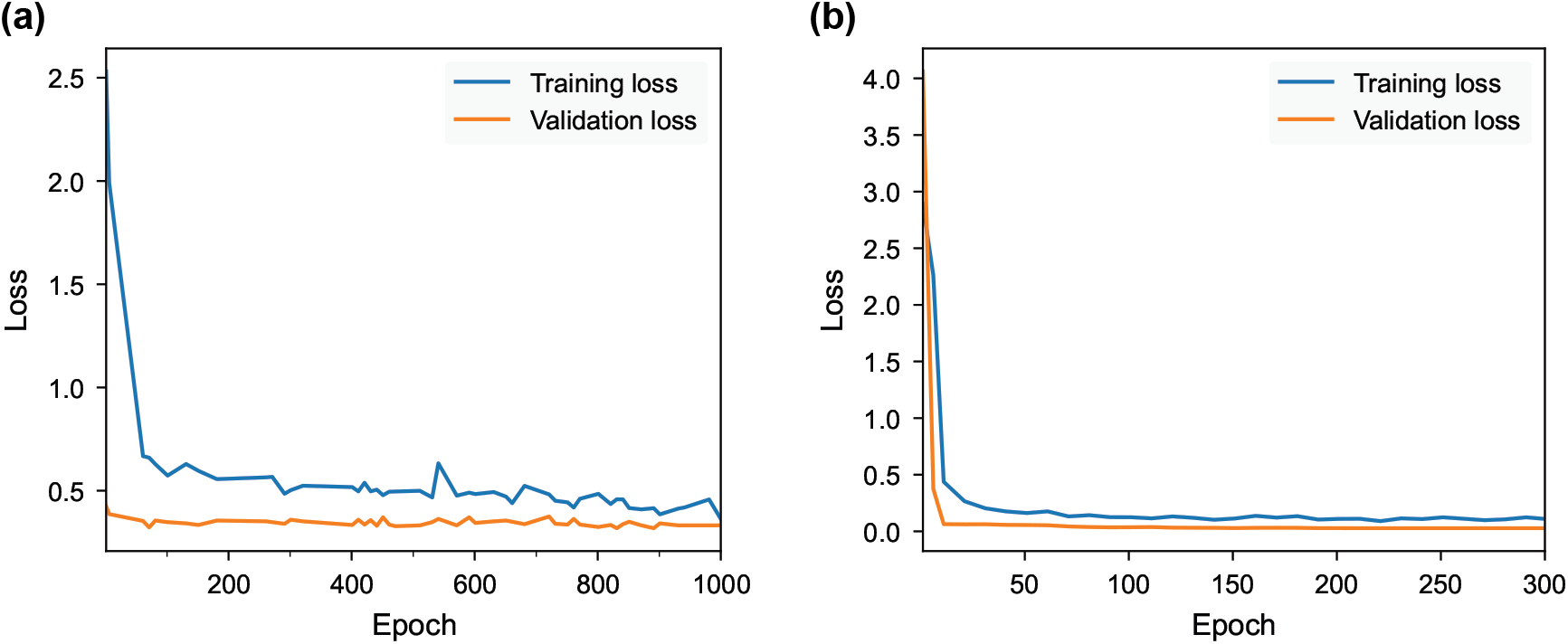
Training and validation losses as a function of epoch of the Cellpose-SAM model fine-tuned for cell (a) and nucleus (b) segmentation.

**Table S1:** Segmentation metrics by target (cells and nuclei) and model (fine-tuned Cellpose-SAM and Amira AI assisted segmentation tool). Aggregated Jaccard Index (AJI) is reported as mean ± SD.

| Target | Model | AJI | $AP_{\tau=0.5}$ |
| --- | --- | --- | --- |
| Cells | Fine-tuned | $0.940 \pm 0.020$ | 0.830 |
| | Amira | $0.916 \pm 0.028$ | 0.818 |
| Nuclei | Fine-tuned | $0.936 \pm 0.043$ | 0.864 |
| | Amira | $0.920 \pm 0.041$ | 0.831 |

**Figure S4:**
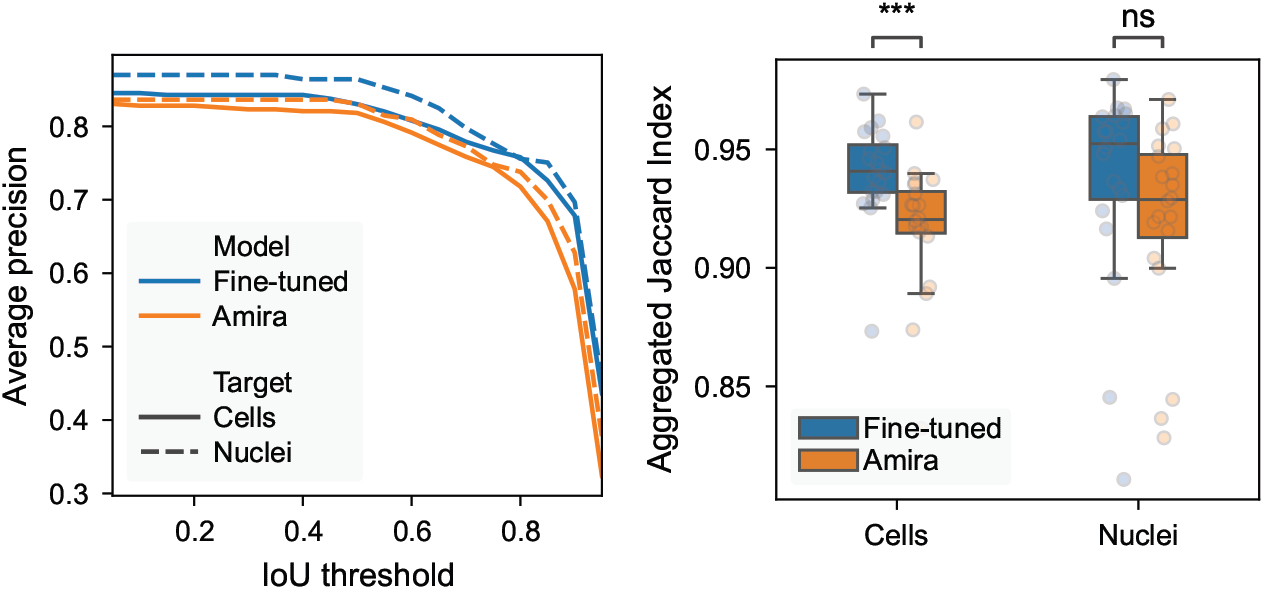
Comparison of segmentation accuracy between fine-tuned Cellpose-SAM and Amira benchmark experiments. Metrics were evaluated on the full labeled dataset (*n* = 20 images). Statistical independence for aggregated Jaccard index populations was evaluated with a Mann-Whitney U test. Significance levels: ***: *p* ≤ 0.001; ns: *p* > 0.05

**Figure S5:**
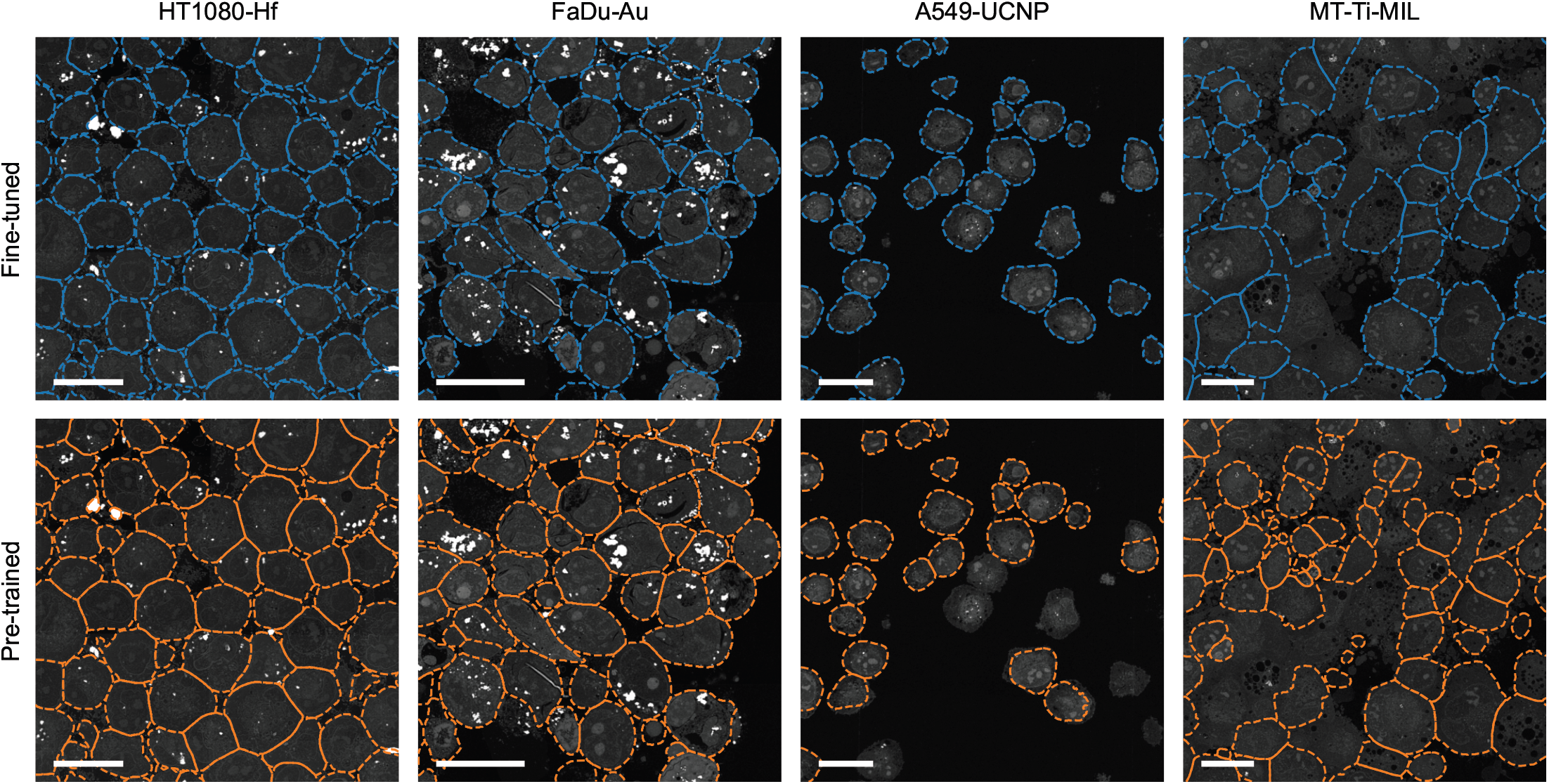
Images of 2D datasets with overlaid cell outlines predicted by fine-tuned (top row) and pre-trained (bottom row) Cellpose-SAM models. Scale bars: 20 µm.

**Figure S6:**
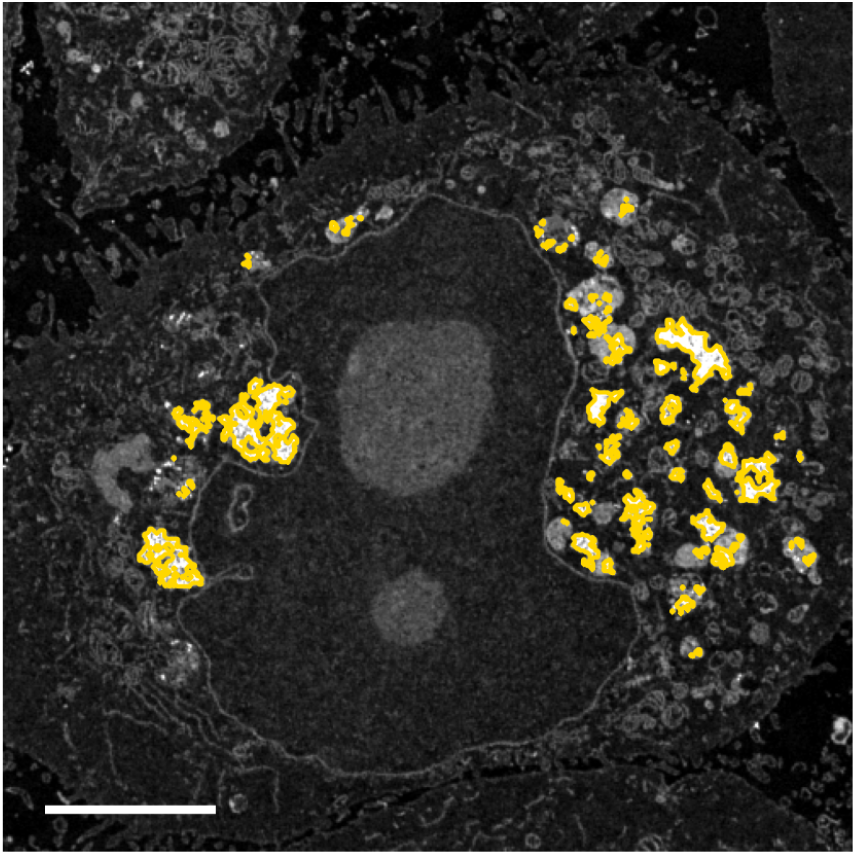
Detail of an image with overlaid NP outlines, calculated by thresholding the NP posterior probability at 95 %. Scale bar: 5 µm.

#### S2.3 Analysis of segmented masks

**Figure S7:**
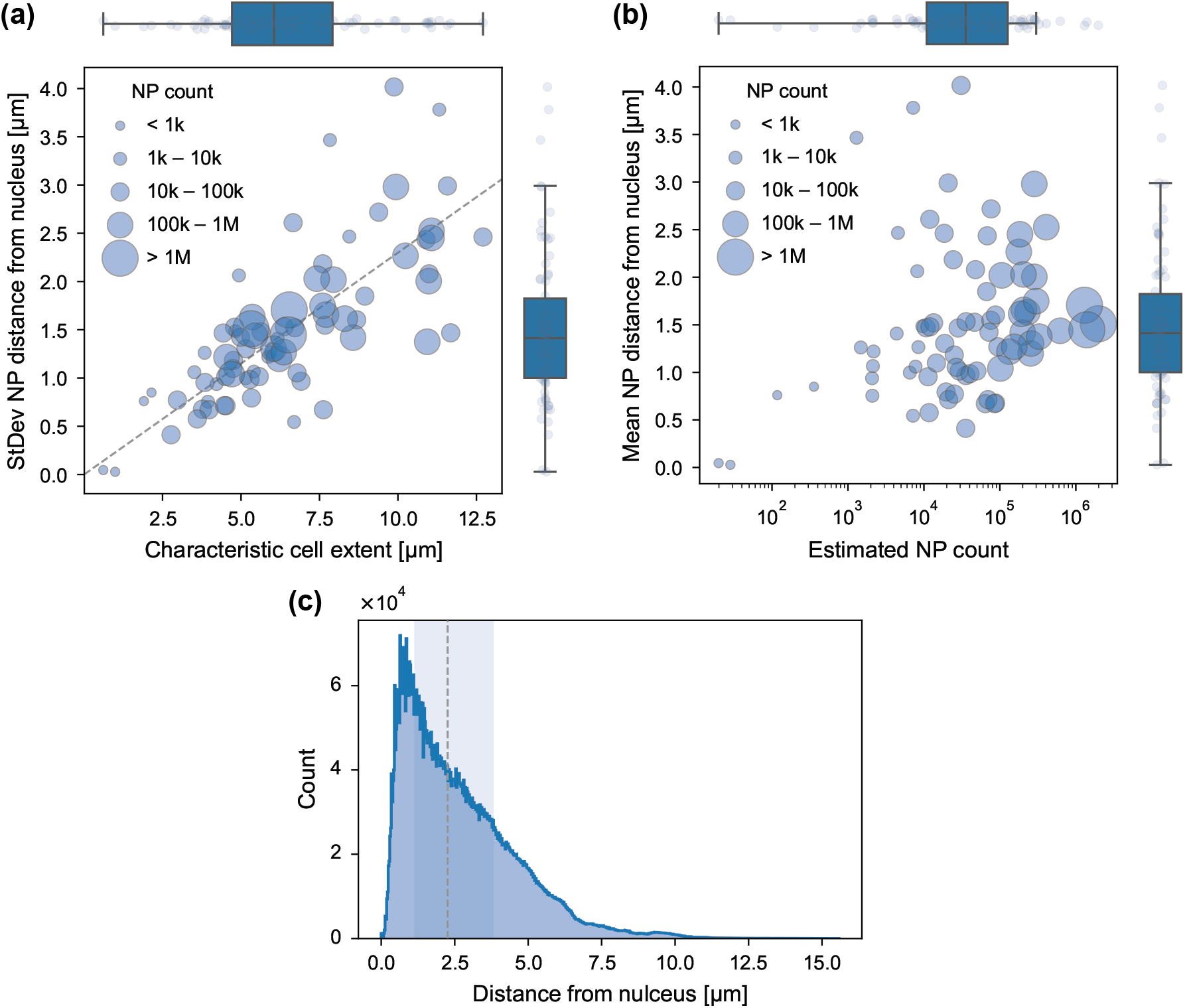
(a): Scatter plot of the standard deviation of distance of NPs from the host cell nucleus as a function of characteristic cell extent (*n* = 78); marker size encodes the estimated NP count internalized per cell; dashed line: linear regression (slope 0.23, intercept 0, *R*^2^ = 0.568). (b): Scatter plot of the mean distance of NPs from the host cell nucleus as a function of the estimated NP count internalized by the cell (*n* = 78). (c): Distribution of distance from the host cell nucleus for all NP voxels (*n* = 7 704 248); dashed line: median; shaded area: Q25–Q75.

